# A quantitative characterization of the heterogeneous response of glioblastoma U-87 MG cell line to temozolomide

**DOI:** 10.1101/2024.05.28.596108

**Authors:** Pragyesh Dixit, Ilyas Djafer-Cherif, Saumil Shah, Arne Traulsen, Bartlomiej Waclaw

**Affiliations:** Dioscuri Centre for Physics and Chemistry of Bacteria, Institute of Physical Chemistry, Polish Academy of Sciences, Kasprzaka 44/52 01-224 Warszawa, Poland; Department of Theoretical Biology, Max Planck Institute for Evolutionary Biology, Plön, Germany; School of Physics and Astronomy, The University of Edinburgh, JCMB, Peter Guthrie Tait Road, Edinburgh, EH9 3FD, United Kingdom

## Abstract

Most cancers are genetically and phenotypically heterogeneous. This includes subpopulations of cells with different levels of sensitivity to chemotherapy, which may lead to treatment failure as the more resistant cells can survive drug treatment and continue to proliferate. While the genetic basis of resistance to many drugs is relatively well characterised, non-genetic factors are much less understood. Here we investigate the role of non-genetic, phenotypic heterogeneity in the response of glioblastoma cancer cells to the drug temozolomide (TMZ) often used to treat this type of cancer. Using a combination of live imaging, machine-learning image analysis and agent-based modelling, we show that even if all cells share the same genetic background, individual cells respond differently to TMZ. We quantitatively characterise this response by measuring the doubling time, lifespan, and motility of cells, and determine how these quantities correlate with each other as well as between the mother and daughter cell. We also show that these responses do not correlate with the cellular level of the enzyme MGMT which has been implicated in the response to TMZ.

## Introduction

Cancer is a disease in which genetic or epigenetic alterations cause cells to proliferate in an uncontrolled way. While many cancers can be attributed to extrinsic factors (smoking, radiation, exposure to UV), more than half are thought to be “bad luck” caused by errors during DNA replication (Tomasetti and Vogelstein 2015; Nowak and Waclaw 2017).

One of the most common modes of cancer treatment, alongside surgery and radiotherapy, is chemotherapy. Traditional (non-targeted) cytotoxic chemotherapy works because cancer cells have a different proliferation or drug uptake rate compared to normal cells, and are thus more susceptible to the drug (Galmarini, Galmarini, and Galmarini 2012). By carefully adjusting the drug dose, cancer cells can be eliminated while minimising the effect on healthy cells. However, the difference between a dose toxic to cancer cells and a dose toxic to normal cells is often small, and even a minor increase in the resistance of cancer cells or the existence of a slightly more resistant subpopulation can lead to treatment failure. Resistance to anticancer drugs can come about in different ways (Luqmani 2008) such as genetic mutations which lead to decreased drug uptake, increased drug efflux, changes in metabolism, or cell death inhibition (Mansoori et al. 2017). Resistance can also be caused by tumour microenvironment – a complex network of interactions between cancer cells and the surrounding normal tissue (Klemm and Joyce 2015; Liotta and Kohn 2001; Tredan et al. 2007). All these mechanisms can lead to substantial heterogeneity in the response of different cells within the tumour to chemotherapy.

Although we now have a very good understanding of genetic heterogeneity in cancer (Gerlinger et al. 2012; Ling et al. 2015; Ryser et al. 2018; Sottoriva et al. 2015; Sun et al. 2017; Xu et al. 2012; Yachida et al. 2010), this is not paralleled by the understanding of non-genetic mechanisms that cause diverse phenotypic responses to drugs. In general, many different mechanisms can cause phenotypic diversity in eukaryotic cells. Two of them, stochasticity in gene expression (Raj and van Oudenaarden 2008) and epigenetic regulation via chromatin modifications (Lund and Lohuizen 2004), are particularly well established. However, their effect on cancer chemotherapy is not entirely clear.

Phenotypic diversity and its relationship to therapeutic response has been investigated in glioblastoma multiforme (GBM), an aggressive brain cancer. GBM is usually treated with surgery, followed by adjuvant radio- and/or chemotherapy. Chemotherapy typically involves alkylating agents such as temozolomide (TMZ) or carmustine (also called BCNU). In particular, TMZ adds methyl groups to guanine in the DNA; an attempt by the cell to repair damaged base pairs sometimes leads to DNA double-strand breaks occurring during cell replication. A large number of methyl adducts thus leads to DNA fragmentation and cell death (Ochs and Kaina 2000). However, cells express the enzyme MGMT (O-6-methylguanine-DNA methyltransferase) which removes methyl groups from guanine (S. Sharma et al. 2009). Early studies demonstrated that MGMT promoter CpG methylation (natural regulatory mechanism, different to TMZ-induced methylation), which reduces the expression of MGMT, correlates with better treatment outcome in TMZ chemotherapy (Esteller et al. 2000). Indeed, MGMT promoter methylation predicts TMZ sensitivity in *in vitro* clonogenic assays (Hermisson et al. 2006). However, it seems that MGMT promoter methylation is not the only factor;(N. R. Parker et al. 2016) have shown that while there is heterogeneity in MGMT promoter methylation in tumours, it does not fully correlate with MGMT expression. They have also detected genetic mutations in other pathways implicated in resistance to TMZ. A recent analysis of single cells harvested from GBM tumours showed the presence of many genetic and epigenetic alterations, and demonstrated that individual clones responded differently to TMZ, which could at least partially be explained by epigenetic alterations in regulatory regions of relevant genes (Akgül et al. 2019).

In this work, we characterise the response of single glioblastoma cells to TMZ. We culture the cells *in vitro*, expose them to TMZ, and optically monitor their behaviour. Taking the advantage of machine-learning approaches to phenotyping (Choi et al. 2021) we track individual cells and perform quantitative analysis of their behaviour. We confirm that there is significant heterogeneity in cell phenotype (division rate, motility, death rate). We then investigate the hypothesis that phenotypic heterogeneity due to non-genetic intracellular variations in MGMT expression can cause different cells to respond differently to TMZ. Using a fluorescently-tagged MGMT as a reporter of MGMT expression we show that there is no correlation between the intracellular concentration of MGMT and cell fate. We conclude that other mechanisms that do not involve MGMT may be more relevant for the observed diversity in the sensitivity of genetically-identical glioblastoma cells to TMZ.

## Results

To investigate the response of glioblastoma cells to temozolomide (TMZ), we used the U-87 MG cell line (Allen et al. 2016). We first performed a clonogenic assay to select an appropriate range of TMZ concentrations for subsequent live-imaging experiments (SI Fig. S1A). The assay showed that the proportion of cells surviving TMZ treatment was close to zero for 500 μM TMZ whereas 10 μM was sufficient to cause a detectable reduction in proliferation. Based on this result, we selected 0, 10, and 500 μM TMZ as the control, low, and high TMZ concentrations. For reference, 10 μM is typical and 50 μM is at the upper end of what has been measured *in vivo* following TMZ chemotherapy, but some *in vitro* studies use concentrations higher than 500 μM (Herbener et al. 2020; Beier et al. 2012).

The design of live-imaging experiments is schematically shown in Figure 1A. We seeded U-87 MG cells in a 96 well plate. After an initial incubation period (16 h), we replaced the cell culture medium with the same medium containing different concentrations of TMZ (0, 10 or 500 μM), and imaged the cells for up to 72 h using a wide-field epi-fluorescent microscope. We then used machine-learning image segmentation to count the number of cells *N*(*t*) at each time point *t*. Figure 1B shows the fold-change *N*(*t*)/*N*(5) (*t* = 5 h chosen as the reference point) for the control (DMSO without TMZ) and TMZ-treated populations. The control population continues to grow at all times, albeit growth slows down in time. In contrast, TMZ-treated populations stop growing after ∼30-40 h of treatment; the fold-change plateaus at different values for low and high TMZ concentrations.

**Fig. 1.**
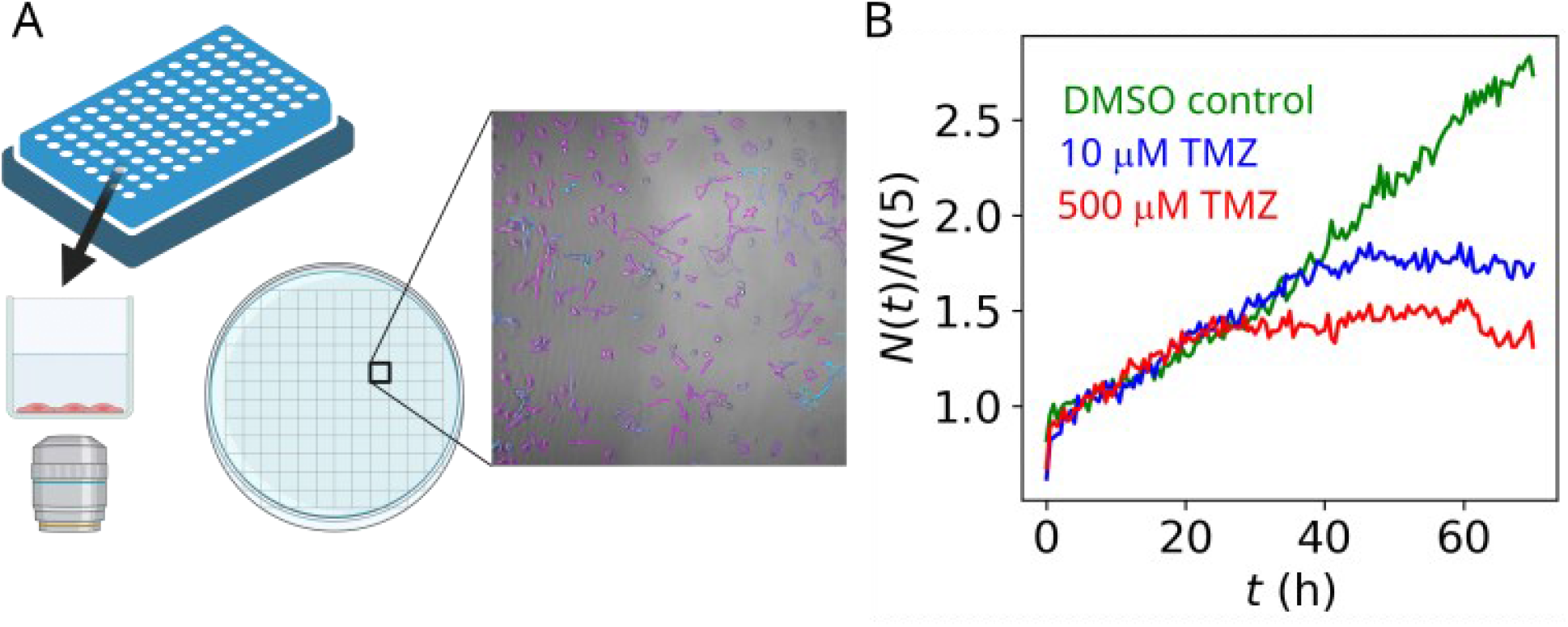
(A) Schematic of the live-imaging experiment. Cells are incubated in the wells of a 96-well plate and imaged through the bottom. Multiple fields of view (FOV) are imaged to cover a large fraction of each well. Imaged cells are detected using a machine-learning segmentation algorithm. (B) Growth curves (fold change as a function of time) of U-87 MG treated with different concentrations of TMZ. The doubling time of the control population is about 40 h. We took *t* = 5 h instead *t* = 0 as the reference cell number *N*(5) due to the non-TMZ related increase in the number of cells for the first few hours (cells settling down after medium exchange).

The cells showed a delayed response to TMZ. Although this has been anticipated based on the known mechanism of action of TMZ (D’Atri et al. 1998), the response was somewhat faster than expected. Previous research (Quiros, Roos, and Kaina 2010) suggested that TMZ should lead to cell death during the 2^nd^ replication cycle since the TMZ exposure. We observed an almost complete cessation of growth after less than one doubling time (Fig. 1B), which may confirm recent suggestions (Strobel et al. 2019) that DNA methylation is not the only mechanism of TMZ-induced cytotoxicity.

A similar behaviour was observed if TMZ-containing medium was replaced with fresh medium after 2 h (SI Fig. S1B), whereas a slightly weaker but very similar response was induced by treating cells with TMZ for only 30 min (SI Fig. S1C). This is consistent with the drug half-life of about 2 h (Strobel et al. 2019; Denny et al. 1994), hence the effective exposure time to TMZ is very short even if the drug is not replaced by fresh medium.

To elucidate the effect of TMZ on individual cells, we tracked the cells and reconstructed cell relatedness from the tracking data (Fig. 2A). Figure 2B shows the number of division events *N*_div_(*t*) versus time, divided by the initial number of cells *N*_0_, for the control and 500 μM TMZ populations. In agreement with population-wide results, we observe that cells continue to divide in the control population; the slope of the *N*_div_(*t*)/*N*_0_ line is 0.026 h^-1^, which is very close to the slope 0.027 h^-1^ of the *N*(*t*)/*N*(5) curve from Fig. 1B. Here we compare the slopes instead of exponential growth rates since growth seems more linear in time than exponential. The TMZ treated cells replicate initially with the same rate as the control cells, but growth slows down significantly after the first 24 h. Since Fig. 2C shows that the death rate is only slightly larger in the treated versus the control population, the reduction in growth observed in Fig. 1B must be predominantly caused by reduced replication rather than increased death.

**Fig. 2.**
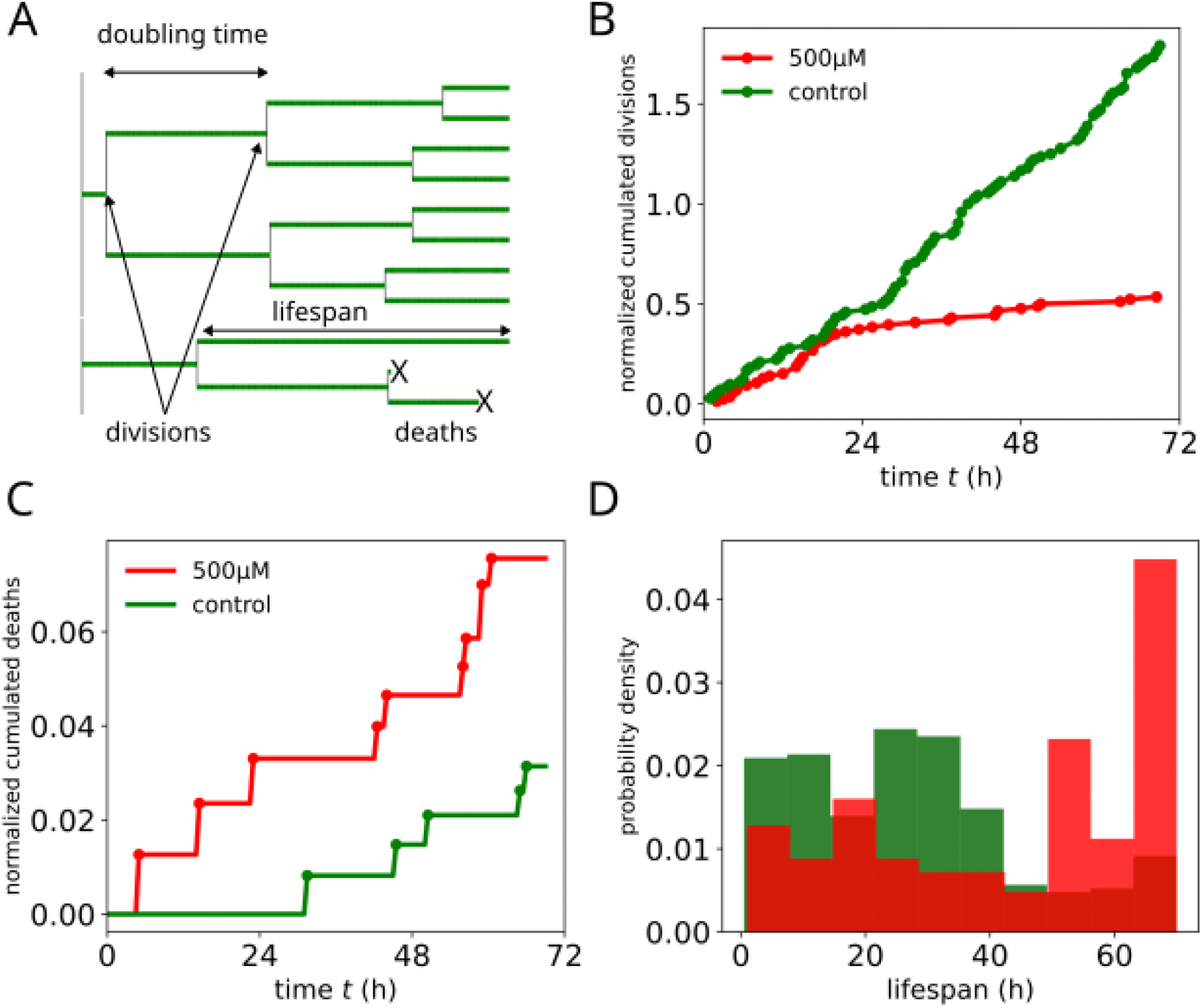
(A) Example of a lineage tree from the control experiment. Arrows indicate two selected divisions and “x” mark two death events; the doubling time is the period of time between two divisions along a single branch of the tree. (B, C) Normalised cumulative divisions *N*_div_(*t*)/*N*_0_ and deaths *N*_death_(*t*)/*N*_0_, where *N*_div_(*t*), *N*_death_(*t*) are the numbers of division/deaths until time *t*, and *N*_0_ is the number of tracked cells at *t* = 0. (D) Probability distribution of cell lifespan for the control (green) and 500 μM TMZ (red).

This shows that TMZ is mostly cytostatic (Günther et al. 2003) during the first days of treatment. However, as the two-week long clonogenic assay shows (SI Fig. S1A), high TMZ concentrations eventually lead to cell death; this is not visible in Fig. 2C due to the experiment lasting only three days.

We also determined cell lifespan, defined as the time since cell’s birth to either death, division, or the end of the experiment, whichever comes first. Figure 2D shows that untreated cells have shorter lifespans compared to treated cells.

Next, we determined the speed of treated and control cells. Figure 3A shows that the average speed of cells in the control population slowly decreases over time. In contrast, the average speed of cells treated with 500 μM TMZ increased sharply after the initial *t* = 12 h (Fig. 3A). At the end of the experiment, TMZ-treated cells were more motile than cells in the control population. Although crowding (increased cell density) could explain lower motility in the control population, less crowding does not explain the sudden increase of motility of treated cells, as their density continued to increase until *t* = 24 h. Figure 3B shows the distribution of mean velocities of cells from the control and 500 μM TMZ populations; it is evident that the proportion of fast-moving cells increased after TMZ treatment. However, if we rescale the two distributions by their average velocities, the distributions are statistically the same (Fig. 3C, *p*_value_ = 0.86). This could mean that treatment increases the motility of each cell by a similar factor as opposed to, e.g. making only a fraction of the population more motile, which would change the shape of the distribution.

**Fig. 3.**
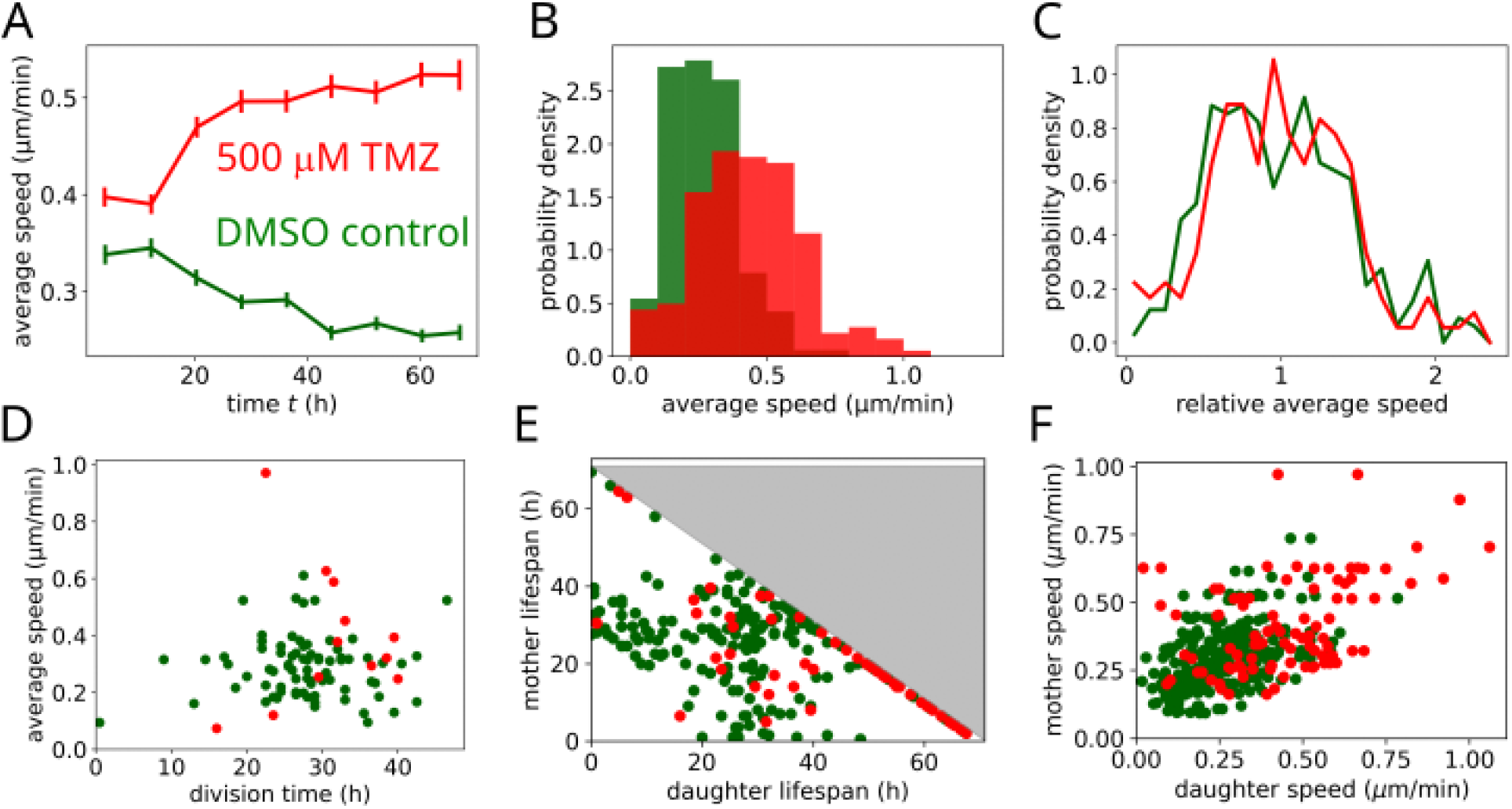
(A) Mean speed (averaged over all tracked cells) as a function of time, for the control and 500 μM TMZ population. (B) Probability distributions of finding a cell with a given average speed in the control and treated populations. (C) The two probability distributions have the same shape when rescaled by the respective mean speed. (D) Average speed versus division time. (E) Mother and daughter cells’ lifespans. Greyed-out area represents lifetimes longer than the duration of the experiment. (F) Correlation between the mother and daughter average velocities. In all plots, green = control, red = 500 μM TMZ.

Those cells that continued to divide showed no correlation between division rates and motility (Fig. 3D), and a small difference between control and treated cells (2D KS test *p*_value_ = 0.008). The lack of proliferation-motility correlation is interesting, as previous research suggested a trade-off between proliferation and motility, i.e. faster-moving cells would be expected to divide less often than slowly-moving cells, or that some sort of division of labour should exist in the population of cancer cells (Giese et al. 2003; Gerlee and Nelander 2012; Evdokimova et al. 2009). We did not find any evidence of such a trade-off in our experiments, though our results are for only a single cell line so it cannot be excluded that other cell lines behave differently. Since Fig. 3D shows only proliferating cells and the control and treated populations have a similar distribution of division rates and velocities, this suggests that treated and non-treated subpopulations of proliferating cells share a similar phenotype. This could mean that cells have different sensitivity to TMZ, and treatment selects a subpopulation of cells (which divide) that are less sensitive.

There is no correlation between the lifespan of mother and daughter cells (Fig. 3E), but we have found a statistically significant correlation between the motility of mother and daughter cells in the control population (Fig. 3F, *R* = 0.46; *p*_value_ < 10^−12^), which suggests that cell phenotype is heritable to some extent. A similar but weaker correlation is visible for TMZ treated cells (Fig. 3F, *R* = 0.46, *p*_value_ < 10^−5^).

These results suggest heritable phenotypic heterogeneity in U-87 MG cells undergoing TMZ treatment. Since, as we indicated in the introduction, MGMT has been implicated in resistance to TMZ, we hypothesised that intracellular variations of MGMT may be responsible for the observed heterogeneous response to TMZ.

To see if varying the intracellular concentration of MGMT affected the response of individual cells to TMZ, U-87 MG cells (approx. 2000/well) were seeded in a 96 well plate and transfected with a plasmid containing a fluorescently tagged MGMT. We imaged the cells for 48 h until some of them started showing bright fluorescence in the nucleus (MGMT exhibits nuclear localization, SI Fig. S2A), and then replaced the medium with a fresh growth medium supplemented with 500 μM TMZ. We continued imaging the cells for another 72 h, tracked individual cells and, using automated image analysis, calculated the total fluorescence in each cell (a proxy for intracellular MGMT concentration). The drawbacks of this approach were (i) not accounting for endogenous, untagged MGMT, (ii) much higher levels of tagged MGMT compared to the WT cells, (iii) MGMT concentration varying in time (SI Fig. S2B). However, since MGMT expression was significantly enhanced in this cell line, we reasoned that any effect of MGMT heterogeneous expression on cell fate should be easy to detect, thus providing a simple test of our hypothesis that MGMT and cell fate are correlated.

Figure 4AB shows cell lifespan (defined above) and motility versus the time-averaged fluorescence of MGMT-C-GFPspark in single cells. We see that, despite a very significant (two orders of magnitude) difference in fluorescence, there is little correlation between motility and fluorescence (control *p*_value_ = 0.56, treated *p*_value_ = 0.63), or lifespan and fluorescence (control *p*_value_ = 0.05, treated *p*_value_ = 0.31). Moreover, this correlation is present in both control and treated cells, and the two data sets are statistically the same (panel A *p*_value_ = 0.17, panel B *p*_value_ = 0.28). This suggests that the correlation is not caused by treatment. A possible explanation is that cells that live longer accumulate more MGMT, so the causation is opposite to what we would expect based on our hypothesis. This means that, at least for the specific cell line used here (U-87 MG), MGMT does not seem to contribute towards resistance against TMZ during the first 72 hours post-TMZ exposure.

**Fig. 4.**
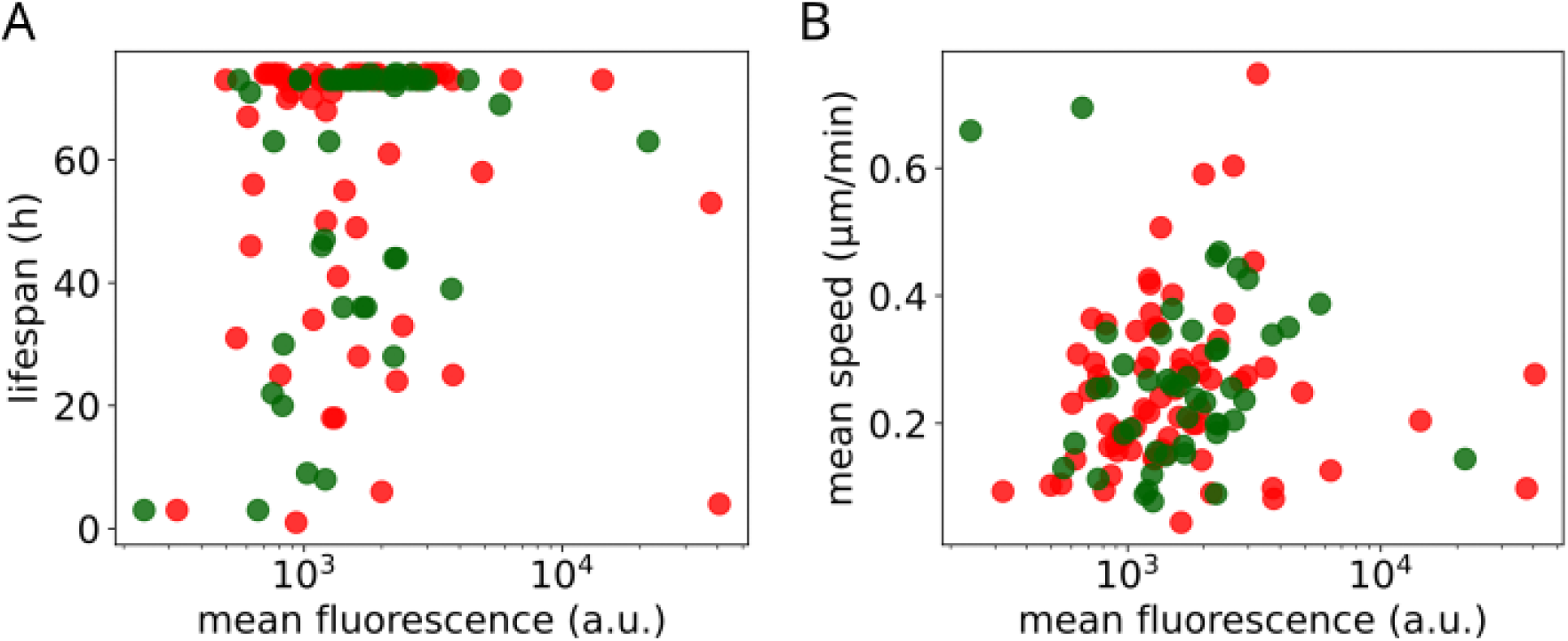
(A) lifespan versus mean fluorescence of MGMT-C-GFPspark, for single cells from the control (green) and treated (red) populations. (B) As in (A) but for mean speed versus mean fluorescence.

### Mathematical model provides insight into the mechanism of TMZ action

To further confirm our understanding of the population dynamics of TMZ-treated cells, we have developed a simple agent-based model that can reproduce the population-level response from Fig. 1B. The model assumes that TMZ methylates some vital components of the cell; the model is agnostic to whether this is DNA or some other molecules such as RNA, histones, or other proteins. Each cell in the model is a separate object with its own internal state, describing the current phase in the cell cycle and whether TMZ-induced methylation has already occurred or not. The cell cycle is divided into three phases; their lengths *T*_1_, *T*_2_, *T*_3_ are the parameters of the model. Cells are methylated only during the second phase with rate *r*_drug_ which depends on the dose of TMZ; *r*_drug_ = 0 when simulating the control experiment. Cells replicate (if not methylated) or die (if methylated) at the end of the cycle. Figure 5A shows the main steps of the computer algorithm used to simulate the model.

**Fig. 5.**
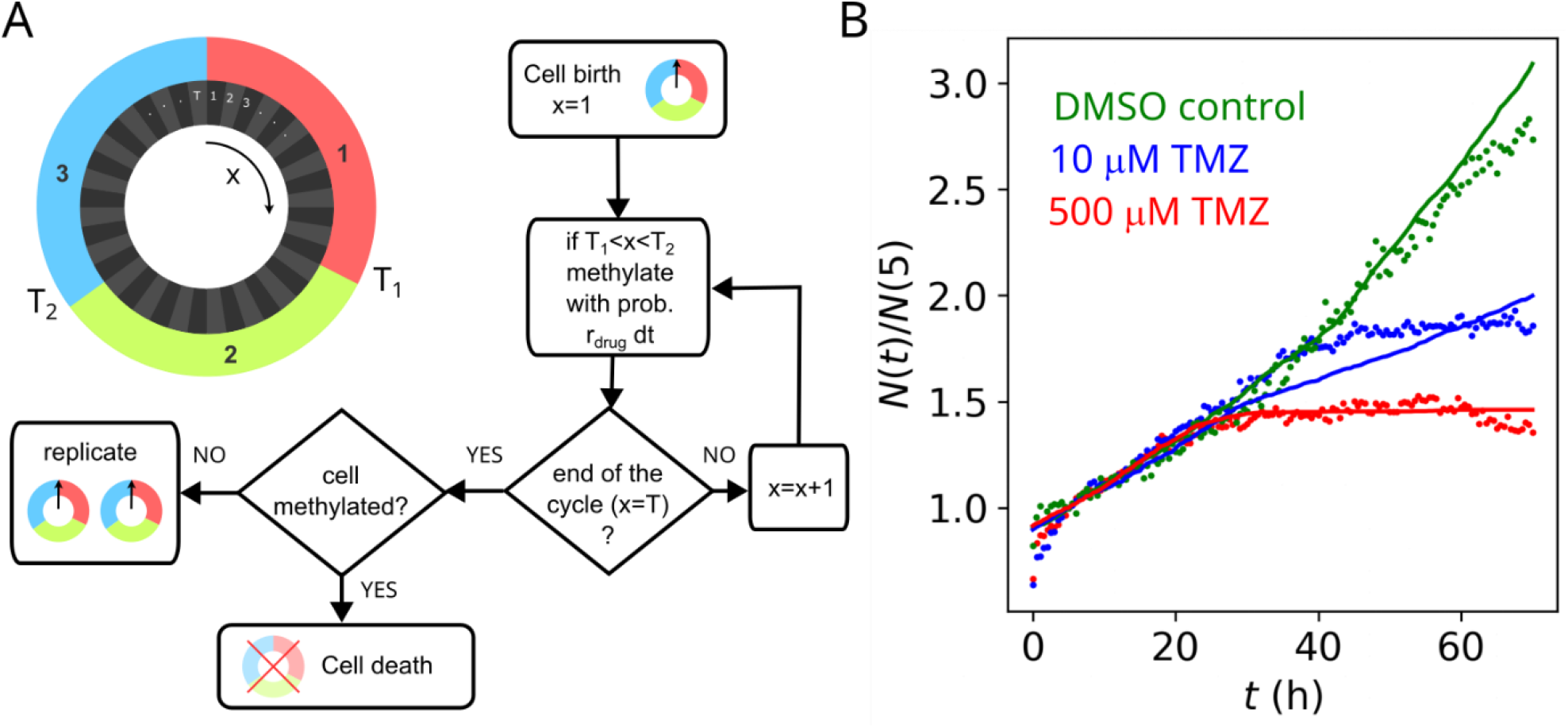
(A) A schematic of the computer algorithm used to simulate the mathematical model. (B) Experimental growth curves (fold-change versus time) compared to the prediction of the model. Points = experimental data (30 min, 2 h and no-TMZ removal data pooled together), lines = computer model (average of 4 replicates per condition).

The model fits the data reasonably well (Fig. 5B). Best-fit parameters are: cell cycle duration *T* = 41 h, methylation starting at the beginning of the cell cycle, *T*_1_ = 0, *T*_2_ = 15 h, and methylation rate *r*_drug_ = 1.83 h^-1^ (500 μM TMZ) and 0.093 h^-1^ (10 μM TMZ). This gives us a potentially interesting insight into the mechanism of action of TMZ. Since the model does not assume that cell death must occur in the 2^nd^ round of replication due to irreparable double-stranded DNA breaks (DSBs) as suggested by earlier works, but it may occur already during the 1^st^ replication round, the agreement between the model and the experimental data supports the idea that TMZ has also non-DNA targets that, if methylated by TMZ, cause cell death much sooner than DSBs. This also agrees with the observed lack of correlation between MGMT level and cell fate, because MGMT is not expected to remove methyl adducts other than those on the DNA. Indeed, there is already evidence (Strobel et al. 2019; Wang, Pickard, and Gallo 2016) that TMZ may have cellular targets other than the DNA.

## Conclusion

In this work we have investigated how glioblastoma multiforme cancer cells responded to *in vitro* chemotherapy with temozolomide (TMZ). We cultured a population of U-87 MG cells in 96 well plates, exposed them to TMZ and imaged for several days using an automated microscope. We initially observed a heterogeneous response to TMZ, i.e. some cells continued to replicate whereas others stopped growing or died. After some time, growth practically ceased but most surviving cells remained motile. Treated cells also become more motile than the untreated control population. Cells that kept dividing exhibited correlations between the phenotype of the mother and daughter cell (similar motility). This complement earlier works on motility of different GBM cell lines in the absence of therapy (De Hauwer et al. 1997) or during targeted therapy (J. J. Parker et al. 2018).

Treated cells slowed down replication after less than a single doubling of the control population. This response was faster than expected based on the suggested mechanism of action of TMZ: DNA methylation followed by failed repair before the first cell division, followed by double strand breaks and death during the second round of replication.

We also performed an experiment in which cells were transfected with a plasmid containing a fluorescently-tagged MGMT protein. The protein was expressed to very different levels in different cells (fluorescence varying by two orders of magnitude) and exhibited nuclear localization typical for the native MGMT. However, we could not find any correlation between the phenotype (motility, doubling time, cell fate) and fluorescence that could be attributed to the action of TMZ.

This, together with the observation that the response to TMZ is faster than expected based on DNA damage-induced cell death, suggests that there must be other mechanisms of action of TMZ that do not rely on DNA damage and can induce a faster response (cell cycle arrest and death). This also means that protection against TMZ does not rely on MGMT alone.

These conclusions are further supported by a mathematical model, which shows that early damage induced by TMZ is sufficient to explain the observed growth curves of control and treated cells.

Our research has several potential shortcomings that could be rectified in future studies:

- No information about the level of endogenous MGMT: we did not fluorescently tag the endogenous MGMT and hence could not determine how diverse its expression was among the cells. However, MGMT expression in U-87 MG is supposedly low (Yi et al. 2019). Moreover, the overexpression experiment did not show any correlation between fluorescently tagged MGMT and the response to TMZ. We would therefore not expect endogenous MGMT to behave differently. This may be different in other cell lines or different concentrations/dosage of TMZ.
- Treatment too short to observe long-term TMZ effects: while the clonogenic assay showed no surviving clones at 500 μM TMZ, only a small fraction of cells died 72 h post-TMZ in live-cell imaging experiments. Longer experiments could be possible but they would be challenging due to the medium being used up, evaporation, and cell crowding.

Regarding the observed lack of correlation between MGMT and the TMZ-induced phenotype, we identified the following potential problems:

- MGMT-C-GFPspark protein may not be active due to misfolding caused by the fluorescent tag. We did not test the enzymatic activity of tagged MGMT. However, others have shown that a similarly fluorescently tagged MGMT exhibited activity towards O6-G methyl groups (Murawska et al. 2022)
- level of overexpressed MGMT so high that it efficiently repairs DNA lesions and the only effect of TMZ we observe is due to non-DNA damage. This would require tagging the native MGMT. We attempted replacing endogenous MGMT with the MGMT-C-GFPspark construct, but could not detect the fluorescent protein in the nucleus using both wide-field and confocal microscopy, despite extensive troubleshooting.

As for the broader significance of our work, phenotypic heterogeneity in the absence of genetic changes has been proposed to be responsible for failure of some anticancer therapies (Brock, Chang, and Huang 2009). Phenotypic diversity may generate resistant cells that can repopulate the tumour when sensitive cells are removed by therapy, or can provide a temporary “safe haven” for cancer cells to evolve genetic resistance – a possibility that has been suggested by others (S. V. Sharma et al. 2010) and independently investigated theoretically by us (Tadrowski, Evans, and Waclaw 2018). However, evolutionary dynamics of non-genetic heterogeneity has not been investigated experimentally in a quantitative way that would enable one to build predictive mathematical models. Here, we attempted to quantify phenotypic heterogeneity in the response to chemotherapy and to determine how it is passed onto offspring cells. While we did not fully succeed in elucidating the mechanism of short-term response to TMZ, our results suggest that MGMT is not involved.

We also obtained quantities such as the distributions of division times, death rate, and cell motility, that could be used to construct more advanced models of glioblastoma treatment. Mathematical modelling is increasingly being used in oncology (Barbolosi et al. 2015; West et al. 2023; Enderling et al. 2019). However, to predict cancer growth and evolution in individual patients, theoretical models must be based on experimentally verified assumptions and carefully measured parameters and their distributions in a population of cancer cells. Most published data cannot be integrated into mathematical models in this way due to different parameters measured for different cell lines, conditions, and experimental approaches. We believe that the quantitative characterization (similar to what we have attempted here) of each patient-derived cancer clone will be required to develop truly predictive models of cancer.

## Materials and Methods

### Cell culture

U-87 MG (ECACC 89081402) cells were cultured in EMEM medium (Corning; catalogue number: 10-009-CV) supplemented with 10% foetal bovine serum (FBS) and 1% penicillin-streptomycin solution. Cells tested negative for mycoplasma contamination (Applied Biological Materials Inc, catalogue number: G238).

In preparation for live-imaging *in vitro* chemotherapy experiments, cells were trypsinized with 1 ml trypsin-EDTA solution for 5 min at 37 °C. Trypsinized cells were centrifuged for 5 min at 300G in EMEM medium. The supernatant was discarded and the cell pellet was resuspended in EMEM medium. 10 μl of the cell suspension was used to estimate the number of cells by an automated cell counter (BioRad TC20). Based on this, 200-500 cells/well were seeded in a 96 well plate in 200 μl of EMEM medium per well. The cells were then left overnight in the incubator (Binder) at 37 °C and 5% CO_2_.

### Drug treatment

After the overnight incubation, the medium was discarded and replaced with fresh EMEM medium without phenol red (Sigma-Aldrich; catalogue number: F0385-500ML), supplemented with 10% FBS, 2 mM L-glutamine (Gibco; catalogue number: A29168-01), 1500 mg/L sodium bicarbonate (Sigma-Aldrich; catalogue number: S8761-100ML), and either 0.25% DMSO (Sigma Life Science; catalogue number: D2650-100ML) or the desired concentration of the drug temozolomide (TMZ, Sigma-Aldrich, catalogue number: T2577-25MG). The concentration of DMSO was adjusted to be the same for all conditions (control and treated populations). Non-DMSO controls were replaced with EMEM without phenol red.

### Clonogenic assay

Cell culture dishes (60 mm) were seeded with approx. 1000 U-87 MG cells/dish in 5 ml EMEM medium. After overnight incubation, medium was replaced with fresh medium containing either 0.25% DMSO or the desired concentration of TMZ. The concentration of DMSO was adjusted to be the same for all conditions. Non-DMSO controls were replenished with EMEM. All the dishes were kept in the incubator for 2 weeks. After the incubation, the medium was replaced with 4% paraformaldehyde for 20 min to fix the cells. Cells were then washed with PBS and stained using 0.5% crystal violet solution for 20 min. After 20 min, cells were washed with water to remove excess crystal violet stain. The dishes were dried at room temperature and the number of colonies formed (visible with a naked eye) were counted manually.

### Expression plasmid and transient transfection

Eukaryotic expression plasmid for human MGMT tagged with GFPspark was purchased from SinoBiological (HG12077-ACG). Fugene HD transfection reagent (Promega; catalogue number: E231A) was used for transient transfections. 100 ng of plasmid DNA was used with the transfection reagent in a 3:1 ratio for 96 well plates. Transfection was done following the manufacturer’s protocol.

### Image acquisition

The 96 well plate with cell cultures was moved from the incubator to the microscope stage. Images were acquired using Nikon Eclipse Ti2-E epi-fluorescent microscope with automated XY stage, Andor Zyla 4.2 SCMOS camera (Oxford Instruments, UK), CO2 incubator (OKO lab), and the Perfect Focus System (Nikon), and controlled by MicroManager 2.0 (Edelstein et al. 2014). While imaging, the plate was incubated at 37°C, with continuous flow of humidified air with 5% CO_2_. Each well was divided into a number of overlapping fields of view (FOV) on a regular grid, with 10% overlap between neighbouring FOVs, covering the entire well. Table 1 shows the specific settings for each experiment. Perfect Focus System was used to adjust for the thermal drift during imaging. Images were analysed using the machine learning algorithms described below.

**Table 1.** Imaging settings for live imaging experiments. BR = brightfield, FITC = green fluorescence filter 457.5 – 487.5 nm (excitation) / 502.5 – 537.5 nm (emission).

| Experiment | Plate type | Magnification | Duration | Imaging frequency | Channels |
| --- | --- | --- | --- | --- | --- |
| Figs. 1,2,3,5 | Corning 3595 | 10 | 72 h | every 30 min | BR |
| Fig. 4 | Cellvis P96-1-N | 20 | 48+72 h | every 1 h | BR, FITC |

### Image and data analysis

All live-imaging experiments generated data in the form of μManager TIFF datasets. We processed these images in two different ways, depending on whether the total count or individual tracks were required:

- *Total count*. Individual bright-field FOVs were segmented using PointRend, a convolutional neural network that segments using point cloud predictions (Kirillov et al. 2020). The algorithm was trained using manually segmented cell images (*n* = 500). The data set was augmented using geometric (scaling, cropping and rotation) and brightness/contrast transformations. Overlapping cells from adjacent FOVs were removed to avoid double counting. The code generated outlines of cells for each time point. The number of such outlines was counted in each time frame; this was used to represent the population size at a given time.
- *Individual tracks*. Individual bright-field FOVs were first stitched using ImageJ Grid/Collection Stitching plugin (Preibisch, Saalfeld, and Tomancak 2009) using a custom-made script for batch processing of all FOVs from all time points. Due to significant changes in cell morphology and a large amount of movement between the frames, automated cell tracking did not work and we resorted to manual tracking using TrackMate 7 (Ershov et al. 2022). A single track represents a collection of (*x, y*) coordinates along the trajectory of a cell, from when it separated from its mother cell or from *t* = 0 (if tracking started at *t* = 0), to the next division event, or its apparent death, or the end of the experiment. In a few cases in which individual tracks apparently merged together, such tracks were excluded from the analysis. Individual tracks were exported as XML files, and imported into a Python notebook using LineageTree library (version 1.3.0). We then segmented individual cells in brightfield images using Segment Anything (Kirillov et al. 2023) using *x, y* and *t* coordinates along the tracks as prompts for the algorithm. These prompts make the algorithm to detect anything that is close to the provided location in the image and forms an object well delineated from the background. We used pretrained weights of the ViT-H model from the package micro-sam, which is a fine-tuned model for microscopy (Archit et al. 2023). This procedure generated binary masks used to determine total fluorescence in each cell (see *Fluorescence of individual cells* below).

#### Growth curves

Fold change in the number of cells was calculated as the ratio of the number of cells *N*(*t*) at time *t* to *N*(*t* = 5), the number of cells after 5 hours from the start of the experiment. We did not normalise by *N*(*t* = 0) because of growth-unrelated increase in the first few hours of the experiment (e.g. cells settling after being disturbed by the addition of TMZ).

#### Motility and division times

For each track, we calculated the instantaneous speed *v*(*t*_*i*_) from the difference in (*x, y*) coordinates at time *t*_*i*_ and *t*_*i*+1_ as *v*(*t*_*i*_) = ((*x*_*i*+1_ − *x*_*i*_)^2^ + (*y*_*i*+1_ − *y*_*i*_)^2^)^1/2^. We then calculated the average speed of each cell as 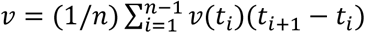, and used this quantity to represent cell motility.

To obtain division times, we calculated the time between the consecutive division events, i.e. *DT* = *t*_*n*_ − *t*_1_ where *t*_1_ and *t*_*n*_ are the first and last points of the track.

#### Fluorescence of individual cells

Uneven image illumination in wide-field fluorescent microscopy causes fluorescence to drop towards the edges of the FOV. To remove this effect, i.e., to apply flat-field correction to the image, we first calculated a “background image” using the rolling ball algorithm applied to stitched fluorescence images. We then divided the original image by the “background image”. Next, we used binary masks of segmented cells to compute total fluorescence of each cell.

#### Statistical analysis

To compare 1D distributions, we used the two-sided Kolmogorov-Smirnov test (Python SciPy 1.10.0). To compare 2D distribution, we used the 2D Kolmogorov–Smirnov test using the code written by Zhaozhou Li, https://github.com/syrte/ndtest (version 0.1) based on Refs. (Peacock 1983; Fasano and Franceschini 1987). Error bars represent standard errors of mean.

#### Mathematical model

We use an agent-based model, in which each cancer cell has four state variables: an integer “clock” *x* = 1, …, *T* representing the current position in the cell cycle, the current cell cycle stage *s* = {1,2,3}, whether the cell is methylated due to the drug *m* = {0,1}, and whether the cell cycle has halted *a* = {0,1}. The parameter *T* represents the duration of the cell cycle as the number of discrete time intervals *dt* = 30 min. We consider three stages in the cell cycle; these could (but do not have to) represent the gap 1 (G1), synthesis (S), and gap 2 (G2) phases. We omit the M stage as the duration of this stage is very short. The phases have durations of *T*_1_, *T*_2_, *T*_3_ time steps, respectively. Thus, the division time is *T* = *T*_1_ + *T*_2_ + *T*_3_ time steps. The value of *x* increases by one every time step of the algorithm. The variable *s* equals 1 for 1 ≤ *x* < *T*_1_, *s* = 2 for *T*_1_ ≤ *x* < *T*_2_ and *s* = 3 for *x* ≥ *T*_2_. When the cell has reached the end of the cell cycle, *x* = *T*, the cell divides into two daughter cells. The two cells start at the beginning of the cell cycle (*x* = 1).

An unmethylated cell (*m* = 0) becomes methylated (*m* = 1) with rate *r*_drug_ (probability per unit of time) during the second phase only. Methylated cells die at the end of the third phase.

The simulation starts with *N*_0_ unmethylated cells, with *N*_0_ taken from the first data point of the experimental data, and with *x* chosen from the exponential distribution *p*(*x*) = (2 ln 2)/*T* 2^−*x*/*T*^. This distribution is the steady-state distribution of *x*, i.e. it represents a population of cells that has been growing for sufficiently long time (as in the experiment prior to the addition of TMZ) for cells to have desynchronized their cell cycles.

#### Fitting the model to data

We fitted the model simultaneously to all growth curves shown in Fig. 5B. All parameters were assumed to be the same for the control, 10, and 500 μM TMZ experiments, except the parameter *r*_drug_ which was permitted to take different values for the three conditions (*r*_drug=0_ = 0 whereas *r*_drug=10_, *r*_drug=500_ were the fitting parameters).

## Data and code

All software used for data analysis (Jupyter notebooks, Julia code), processed data and simulation results are available at https://github.com/Dioscuri-Centre/phenotypic_heterogeneity. Due to large file sizes (several TBs), access to raw image data will be provided on request.

## Use of AI tools

No AI tools were used for manuscript write-up and editing. Convolutional neural networks and vision transformers were used for image processing (Methods).

