## Supplementary Figures for "A quantitative characterization of the heterogeneous response of glioblastoma U-87 MG cell line to temozolomide"

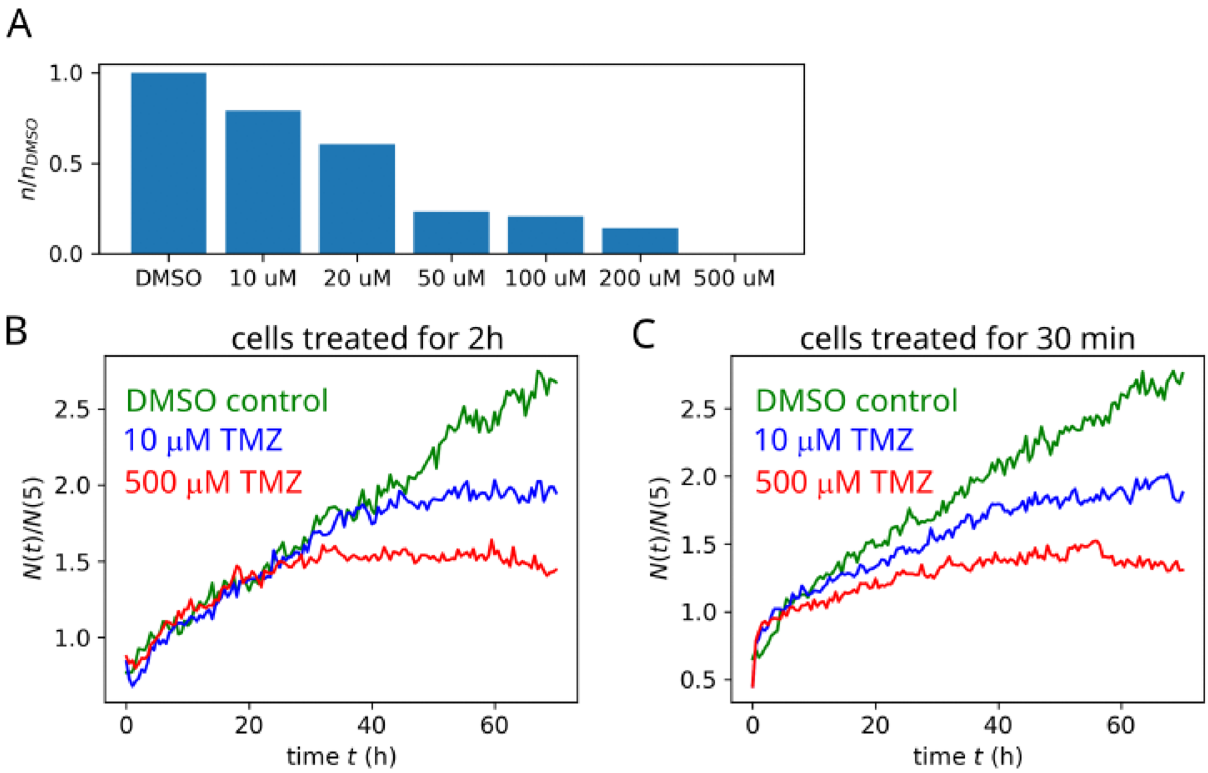


**SI Fig. S1.** (A) Clonogenic assay: the ratio $n/n_{\mathrm{DMSO}}$ between the number of colonies $n$ from the TMZ-treated cultures to the number of colonies $n_{\mathrm{DMSO}}$ in the control population (DMSO only) formed after 2 weeks of incubation. (B, C) Fold-change in the number of cells versus time for cells treated with TMZ for 2h (B) and 30 min (C).

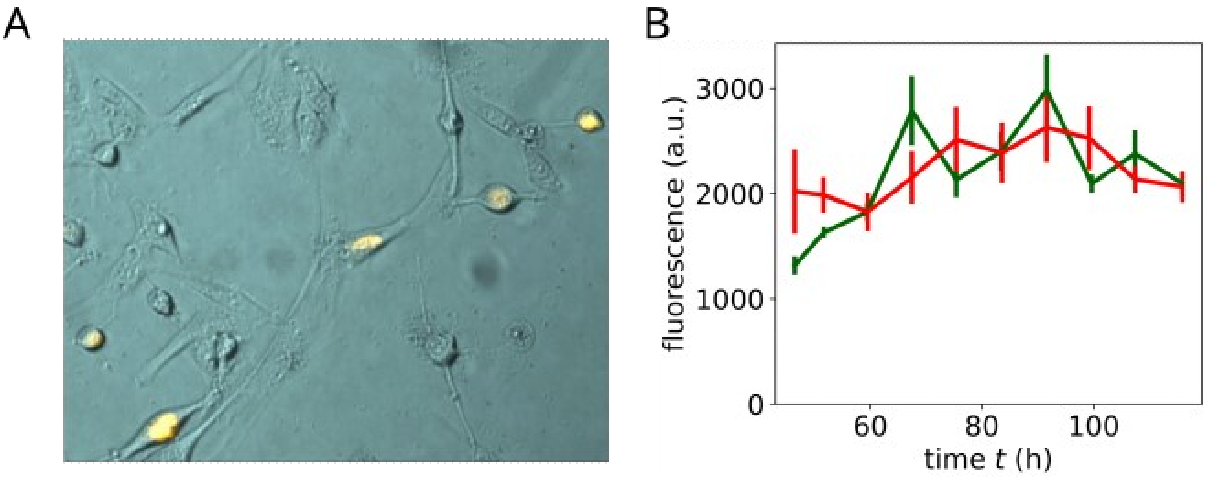


**SI Fig. S2.** (A) Nuclear localization of MGMT-C-GFPspark. Fluorescence (yellow) is superimposed on a brightfield image (teal). (B) Fluorescence versus time for the control (green) and 500 μM TMZ (red), starting from day 2 post-transfection.
